# Early-onset spontaneous murine mammary tumors show increased lymphocyte representation and reduced MHC antigen presentation

**DOI:** 10.64898/2026.10.12.758076

**Authors:** Ryann Ray, Lisa Polin, Rui Geng, Benjamin Kidder, Jennifer Jacob

## Abstract

**Simple Summary:** Not all breast tumors behave the same way, and part of the reason may be inherited rather than caused by the tumor itself. Using mice that carry the same cancer-causing HER2/neu gene but differ in genetic background, we found that introduction of the PWK/PhJ genetic background was associated with tumors that appeared about a month earlier. These early tumors contained a larger proportion of immune cells, including T and B cells, but the T-cell population showed reduced expression of genes associated with cytotoxic activity. At the same time, the tumors had lower levels of proteins and genes that let the immune system recognize tumor cells, alongside higher levels of structural proteins that can make tissue stiffer and easier for cancer to invade. These findings suggest that genetic background shapes not only how quickly breast tumors arise but also how well the immune system can recognize them, a distinction that may matter for future immune-based treatments.

**Background/Objectives:** Genetic background modifies cancer risk and phenotype in humans, but its influence on the tumor immune microenvironment and antigen-presentation machinery in ErbB2 (HER2/neu)-driven breast cancer is not well defined. We used a genetically diverse mouse cross to test whether the wild-derived PWK/PhJ background alters immune infiltration and antigen presentation in a NeuT model of HER2/neu-positive breast cancer.

**Methods:** Previously, quantitative trait locus (QTL) mapping using Diversity Outbred mice and r/qtl2 identified candidate loci driving tumor onset and growth. The wild-derived PWK/PhJ genetic background was predicted to harbor genes that regulated early onset of HER2/neu-driven tumors. BALB NeuT and (BALBxPWK) F1 NeuT female mice were monitored twice weekly for spontaneous mammary tumor onset and growth. Tumor immune composition was profiled by single-cell RNA sequencing. Protein abundance was measured by data-independent acquisition proteomics in formalin-fixed, paraffin-embedded tumor tissue (n=5 per genotype). Single-nucleus multiome (RNA + ATAC), bulk RNA-seq, and H3K27ac datasets from the same strain comparison were analyzed for focused MHC II cross-layer analyses.

**Results:** As predicted, (BALBxPWK) F1 NeuT mice developed tumors significantly earlier than the parental BALB NeuT mice (10 vs. 14–15 weeks of age; *p* < 0.0001). In the scRNA-seq dataset, F1 tumors showed greater representation of *Cd3e*-expressing T cells and a *Cd79a*-expressing B-cell population not detected in BALB NeuT tumors, and a distinct neutrophil cluster; F1 T cells also showed lower expression of the cytotoxicity-associated genes *Tox, Ltk, Nkg7*, and *Gzmb* relative to BALB NeuT tumor-infiltrating T cells. Proteomics identified 46 differentially abundant proteins (25 higher, 21 lower in F1), with coordinate down-regulation of *Stat1, Tapbp, Parp9*, and *Dtx3l* alongside MHC class I and II proteins and up-regulation of extracellular-matrix and elastic-fiber proteins. Focused cross-layer analysis showed lower whole-tumor expression of a *Stat1*/MHC II-associated gene set together with reduced epithelial chromatin accessibility at selected regions near *Cd74* and *H2-Aa*.

**Conclusions:** The PWK/PhJ genetic background is associated with earlier HER2-driven tumor onset, greater lymphocyte representation, and reduced activity of a STAT1-associated antigen-presentation program despite similar post-onset tumor growth kinetics.

## INTRODUCTION

Human epidermal growth factor receptor 2, HER2, is a transmembrane receptor with tyrosine kinase activity that drives cell growth, proliferation and survival signals^1^. Overexpression or amplification of HER2, encoded by ERBB2, thus defines an aggressive subset of breast cancers^2–4^. The BALB-NeuT mouse model expresses a constitutively active, mutated version of the rat ErbB2 oncogene called NeuT^5, 6^. NeuT is expressed under the mouse mammary tumor virus promoter, faithfully recapitulates multifocal, spontaneous mammary tumorigenesis and has been used to dissect mechanisms of oncogene-driven tumor progression and to test preventive and therapeutic interventions^7–10^.

Inbred BALB NeuT mice develop mammary tumors on a fixed, largely invariant genetic background, which is not representative of outbred humans. Previously, we crossed BALB NeuT males with Diversity Outbred (DO) females to generate (BALBxDO) F1 NeuT mice. DO mice are a genetically heterogeneous outbred strain that captures the allelic diversity of eight founder strains, including the wild-derived PWK/PhJ strain. (BALBxDO) F1 NeuT female mice developed spontaneous mammary tumors with a broad range of tumor onset ages and growth rates^11^. Haplotype reconstruction and quantitative trait locus (QTL) mapping using r/qtl2^12, 13^ identified QTL on chromosomes 1 and X that associated with tumor onset age, and a QTL on chromosome 10 that was associated with tumor growth rate. The PWK/PhJ haplotype showed consistent associations with earlier tumor onset and faster tumor growth across the identified QTL regions (**Figure 1**).

**Figure 1.**
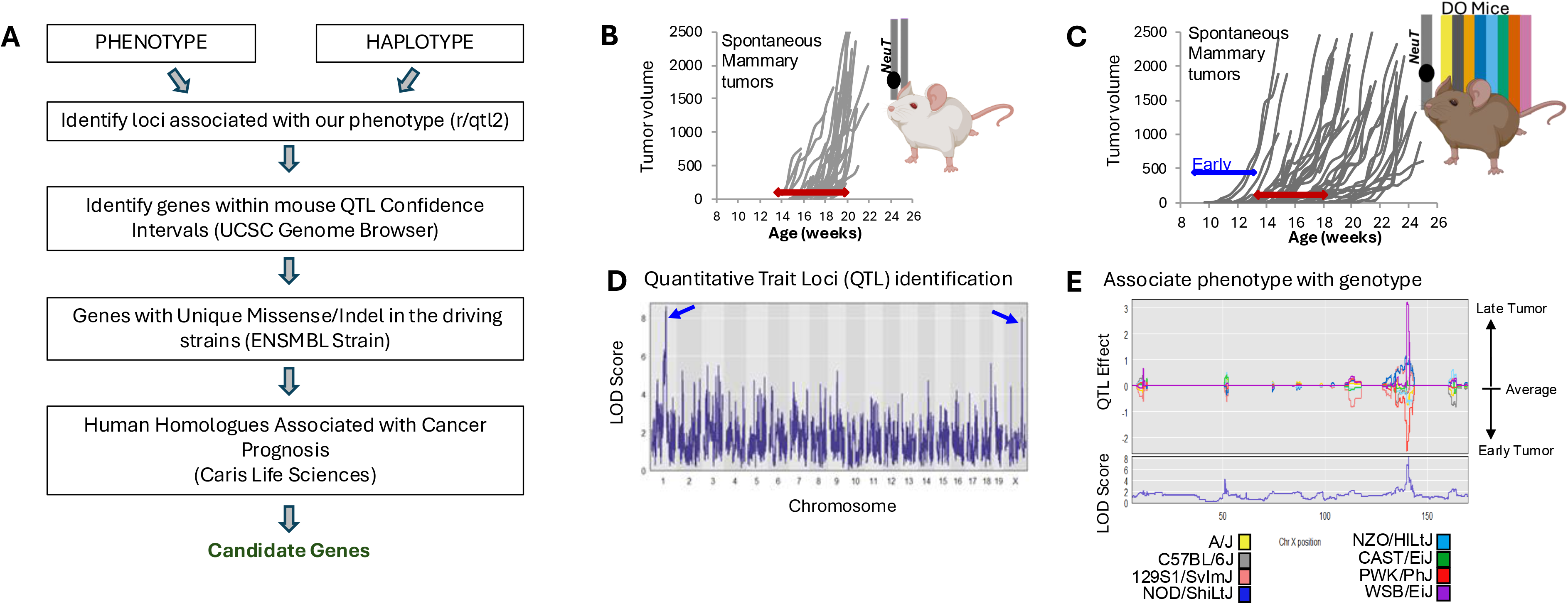
Genetically heterogeneous mice identified the PWK background as driving early tumor onset in a model of HER2/neu-driven breast cancer. (A) The workflow used to identify candidate genes involved in tumor onset and growth. The BALB NeuT model is a murine model of breast cancer on the BALB/c background that expresses the rat homologue of human ERBB2 (called neu) and drives spontaneous mammary tumorigenesis. The phenotype used for this correlation was the age (in weeks) when the total tumor volume in any or all mammary glands exceeded 50 mm^3^. Haplotypes of each mouse was generated by running GigaMUGA SNP arrays on genomic DNA. The phenotype for each mouse was associated the haplotype using the r/qtl package, possible QTL peaks identified across the chromosomes, and the founding strain(s) driving the phenotype identified. (B) NeuT-driven spontaneous mammary tumors become palpable between 14 and 19 weeks of age in the BALB/c background and grow progressively. BALB NeuT mice carry one BALB/c chromosome and the other carries the NeuT transgene (*gray bars*). (C) BALB NeuT male mice were crossed with Diversity Outbred (DO) females and the resulting neu-expressing F1 females monitored for tumor development. The F1 offspring carried the NeuT transgene in the chromosome from the BALB NeuT male (gray bar) and a mosaic chromosome made up of the 8 DO backgrounds (colored bars). (BALBxDO) F1 NeuT female mice developed tumors with significantly higher variability, both in tumor onset age and growth rates. *Red bars* indicate onset age in the BALB NeuT mice. A set of mice developed tumors significantly earlier than the remaining mice and are highlighted by the *blue bar*. (D) A Manhattan plot identifying QTL in murine chromosomes 1 and X when the phenotype of onset age was used. Chromosome 10 was also predicted to harbor a QTL when tumor growth rate was analyzed^11^. (E) The strain effect plot identifies which strain(s) from the DO are correlated with the phenotype in QTL regions. This example QTL on Chr X shows that the PWK/PhJ haplotype is associated with earlier tumor onset. [mouse schematics generated using BioRender]

Tumor cells will adapt a number of mechanisms to avoid destruction by the immune system^14–16^, including down-regulation of the antigen-presentation pathway. The tumor immune microenvironment and the integrity of antigen-presentation machinery are critical determinants of tumor progression^17^ and response to HER2-directed and immune-based therapies^18–21^. Major histocompatibility complex (MHC) class I molecules present endogenous peptides to CD8+ T cells, while MHC II molecules present exogenous peptides. Loss of MHC class I surface expression or of components of the antigen-processing machinery (e.g., TAP1, TAP2, tapasin/TAPBP, ERAP1) is one of the most common and best-characterized mechanisms by which tumors escape T cell-mediated immune surveillance, rendering cells invisible to cytotoxic T cells even when they harbor immunogenic mutations^22, 23^. An MHC-I-low phenotype has been described across many common human cancers, including breast cancer^24, 25^, and MHC class I and TAP1/TAP2 down-regulation was reported more than two decades ago in high-grade primary breast carcinoma lesions^26–28^.

Loss of antigen-presentation also has clinical consequences in breast cancer. In triple-negative breast cancer (TNBC), reduced tumor expression of MHC class I antigen-presentation pathway proteins — including antigen presentation machinery such as TAP1, TAP2, calreticulin, HLA-A, ERAP1, and TAPBP — is associated with significantly shorter recurrence-free and overall survival^29^. Tumor-cell expression of MHC class II, typically not constitutively expressed by most epithelial cells, can be induced by interferon-γ signaling^30–33^, and correlates with tumor-infiltrating lymphocyte abundance, tertiary lymphoid structure formation, and improved disease-free survival in node-positive TNBC^34–36^. In HER2-driven disease specifically, HER2/ERBB2 signaling itself can be inversely correlated with MHC class I expression and impair T cell recognition of peptide:MHC-I complexes^37, 38^, while mechanistic work has identified regulators such as NLRC5 and the transmembrane protein MAL2 that control MHC-I surface abundance and antigen presentation in breast tumor cells^39–42^. Antigen-presentation loss is thus mechanistically and clinically linked to both interferon signaling and HER2 pathway activity.

Whether the same germline genetic background that accelerates ErbB2-driven tumor onset also reshapes tumor immune cell composition and antigen-presentation programs has not been examined. Here, using single-cell RNA sequencing and quantitative proteomics, together with focused analysis of multiome, bulk RNA-seq, and H3K27ac datasets, we show that (BALBxPWK) F1 NeuT tumors develop earlier than BALB NeuT tumors and, in the scRNA-seq dataset, show greater representation of CD3+ T cells and a distinct neutrophil population, together with coordinate down-regulation of MHC antigen-presentation and interferon-stimulated proteins and up-regulation of extracellular matrix (ECM) remodeling proteins.

## MATERIALS AND METHODS

### Tumor growth in BALB NeuT mice

All procedures were performed in accordance with the US Public Health Service Policy on the Use of Laboratory Animals and with approval by Michigan State University’s Institutional Animal Care and Use Committee.

BALB NeuT mice (provided by G. Forni and F. Cavallo, University of Torino, Torino, Italy) were maintained in our facility by mating male NeuT mice with purchased 6-8 week old female BALB/c mice^9, 10^. Mice were maintained in standard animal housing. Homozygous Neu^+/+^ mice die *in utero*, thus the presence of the remaining copy of the NeuT transgene was determined with PCR^11^ from 0.5 cm of tail tissue when pups were 21 days of age^43, 44^. We collaborated with Jackson Laboratories to mate female PWK/PhJ mice with heterozygous male BALB NeuT mice at the Jackson Laboratory to generate (BALBxPWK) F1 NeuT mice.

Female progeny were monitored for onset of mammary tumors by palpation beginning at 5 weeks of age. Tumor volume was calculated as mm^3^ = (X^2^xY)/2, where X is the shorter side and Y is the longer side, then all tumor volumes were summed across the 10 mammary glands. Date of tumor onset was the date at which the total tumor volume was greater than 50 mm^3^.

### Single-cell RNA sequencing

Spontaneous tumors from 65 to 300 mm^3^ without visible necrosis were dissected out from surrounding mammary fat pad tissue. Tissue was minced finely with scalpels and enzymatically dissociated (2.5 mg/mL Collagenase D [Millipore Sigma, Burlington, MA], 5 mg/mL Collagenase type 2 [Worthington-Biochemical, Lakewood, NJ], and 60 U/mL DNAse I [Sigma Aldrich] in RPMI media)^45^ by incubating, shaking, at 37 °C for 30 minutes. EDTA (0.5 M) was added and samples were incubated at 37 °C for an additional 5 minutes. Following cell straining through a 70 μm cell filter, RPMI with 10% FBS (R10) was added to q.s. to 40 mL and centrifuged at 500 x *g* for 10 minutes with reduced deceleration to assist in separating adipocytes. Cell pellets were washed thrice with R10 and centrifuged as before. Cells were resuspended and red blood cells lysed by addition of ammonium-chloride based RBC lysis buffer (ThermoFisher) for 5 minutes. Lysis was stopped by addition of R10 and cell suspensions were filtered through a 40 μm cell strainer and centrifuged as before. Cell pellets were resuspended and viable cells concentrated using the MojoSort magnet bead dead cell removal kit (BioLegend). Samples with viability above ∼90% proceeded to single cell library construction using the 10X Genomics Chromium Next GEM Single Cell 3’ Reagent kit (v3.1 chemistry) and Chromium Single Cell Controller per the manufacturer’s instructions (https://assets.ctfassets.net/an68im79xiti/1eX2FPdpeCgnCJtw4fj9Hx/7cb84edaa9eca04b607f9193162994de/CG000204_ChromiumNextGEMSingleCell3_v3.1_Rev_D)^46^. cDNA libraries were sequenced (NextSeq 500, Illumina, San Diego, CA) to a saturation of 60-70%, with an average of approximately 50,000 reads per cell at the Center for Molecular Medicine and Genetics Genomics Core, Wayne State University or Genomics Core, Michigan State University. Reads were demultiplexed, aligned to the refdata-cellranger-mm10-2020-A, barcodes processed and UMI counts obtained using 10X Genomics’ Cell Ranger version 4.0.0. The transcript profiles of cell clusters were integrated and analyzed using the R-based package Seurat v4.0.6 (https://satijalab.org/seurat/index.html) using UMAP dimensional reduction with a clustering resolution of 0.3^47^. t-SNE visualizations for **Figure 3** were generated using the 10X Genomics’ Loupe Cell Browser. Cluster identities were assigned from top marker genes using Enrichr^48^ and checked against Tabula Muris^49^.

### Proteomics analysis

Tumors from BALB NeuT and (BALBxPWK) F1 NeuT female mice (n=5 each) with tumor ∼300 mm^3^ and without visible necrosis were collected into 10% formalin and embedded in paraffin (FFPE). 20 μm thick tissue sections were removed and sent for Data Independent Acquisition (DIA) in collaboration with the IDeA National Resource for Quantitative Proteomics (Little Rock, AR). Following FFPE tissue processing and tryptic digestion, desalted peptides in aqueous solution were injected for data acquisition from 96-well plates. Up to 1 µg of each peptide sample was drawn into a 10 μL sample loop by the UltiMate 3000 autosampler and loaded onto a 0.5 cm x 75 μm trapping column (Waters XSelect CSH C18 3.5 μm resin). Peptides were then separated by an acetonitrile gradient on a 15 cm x 75 μm analytical column (Waters XSelect CSH C18 2.5 μm resin) over a 60 min data independent acquisition. The Exploris 480 is configured to acquire a precursor scan at 60,000 resolution, followed by 50 × 12 m/z DIA spectra (12 m/z precursor isolation windows at 15,000 resolution) using a staggered window pattern with optimized window placements. Data was searched using Spectronaut against the UniProtKB mouse specific database using the directDIA method with an identification precursor and protein q-value cutoff of 1%, generated decoys set to true, protein inference workflow set to maxLFQ, inference algorithm set to IDPicker, quantity level set to MS2, cross-run normalization set to false, and protein grouping quantification set to median peptide and precursor quantity. MS2 intensity values was assessed for quality using ProteiNorm. Data were normalized using the method that provided the least variability among sample group replicates and highest replicate correlation, and then analyzed using proteoDA to perform statistical analysis using limma with empirical Bayes smoothing to the standard errors. Proteins with an FDR adjusted *p*-value <0.05 and a fold change >2 was considered significant. Data files were processed through EncyclopeDIA or Spectronaut database search. Identified proteins were assessed for abundance between both genetic backgrounds.

### Multiome RNA/ATAC and H3K27ac analysis of MHC II and antigen presentation

For the focused analysis in Figure 5, we analyzed single-cell multiome data generated for the companion PWK NeuT genomics study, comprising three BALB NeuT and three (BALBxPWK) F1 NeuT tumors^50^. Curated weighted-nearest-neighbor annotations were used, retaining tumor epithelial and luminal-alveolar states. MHC II module scores were calculated from normalized RNA expression of *Ciita, Cd74, H2-Aa, H2-Ab1, H2-DMa, H2-DMb1*, and *Ctss* after gene-wise scaling across retained epithelial cells. Scores were averaged within tumor-state combinations containing at least 10 cells, then across eligible tumors with equal weighting within each strain. State-level strain differences were projected onto the epithelial UMAP for descriptive visualization. Epithelial pseudobulk RNA counts were summed within each tumor, filtered using edgeR, normalized using the trimmed mean of M-values method, and analyzed using quasi-likelihood negative-binomial models with strain as the predictor. Benjamini-Hochberg false-discovery rates (FDRs) were calculated across tested genes. Whole-tumor RNA effects and FDRs were obtained from a previously generated bulk RNA-seq edgeR analysis from the same strain comparison^50^. Duplicate gene annotations were represented by the lowest-FDR entry for visualization. Log₂ fold changes represented (BALBxPWK) F1 NeuT relative to BALB NeuT.

The seven module genes plus Stat1 and Irf1 were evaluated using an exploratory one-sided Wilcoxon rank-sum test comparing their whole-tumor log₂ fold changes with those of all remaining genes with finite estimates. The alternative hypothesis was that the selected genes had lower (BALBxPWK) F1 NeuT-relative-to-BALB NeuT effects. Epithelial ATAC counts were summed across the same retained populations within each tumor and analyzed using the edgeR quasi-likelihood workflow. Candidate peaks for Cd74 and H2-Aa were selected within 20 kb of annotated mm39/GRCm39 transcription start sites. Gene-associated bulk H3K27ac intervals from the companion genomics dataset were obtained from the integrated region-to-gene table, retaining the lowest-FDR interval per gene. These selected interval effects were integrated with whole-tumor and epithelial RNA effects for cross-layer comparison. Differential RNA and chromatin analyses used biological tumors as replicates. Cross-layer integration compared strain-level effect estimates and was not treated as a paired multi-assay analysis of the same individual tumors.

## RESULTS

To test whether the predicted PWK genetic background was associated with altered tumor onset, we generated the (BALBxPWK) F1 NeuT cross and found that F1 females developed tumors substantially earlier than parental BALB NeuT mice (**Figure 2**). F1 mice developed palpable tumors by 10 weeks of age (**Figure 2A**) and continued to grow progressively (**Figure 2B**). Tumor-free survival differed significantly between groups (**Figure 2B**). Although tumor onset occurred earlier in (BALBxPWK) F1 NeuT females, growth rates calculated as the days to 500 mm^3^ were not significantly different (8.7<u>±</u> 5.3 days in F1 versus 9.4<u>±</u> 4.7 days in BALB NeuT).

**Figure 2.**
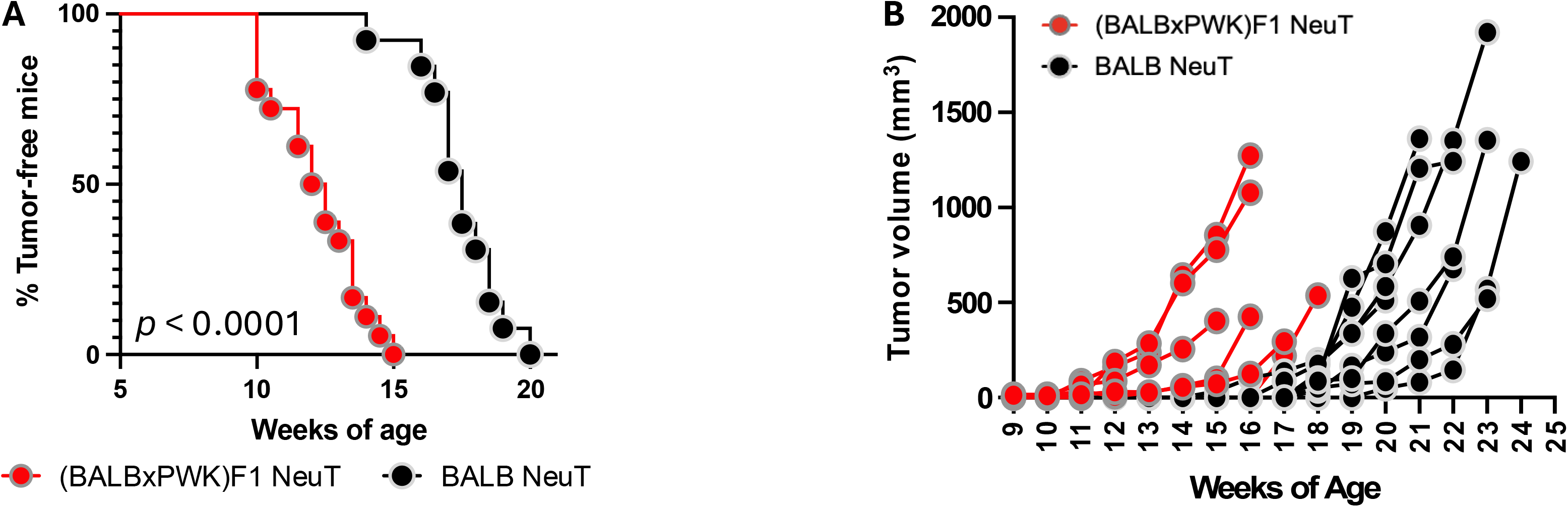
(BALBxPWK) F1 NeuT female mice develop spontaneous mammary tumors significantly earlier than BALB NeuT mice. BALB NeuT and (BALBxPWK) F1 NeuT female mice develop spontaneous mammary tumors and were thus monitored for tumor development by twice-weekly palpation beginning at 5 weeks of age. **(A)** (BALBxPWK)F1 NeuT (*red*) developed tumors which grew progressively beginning at week 10. BALB NeuT (*black*) mice developed tumors beginning at 14 weeks of age, that also grew progressively. Tumor volume is calculated by the summation of all mammary tumors in each mouse and the total tumor volume is shown. **(B)** A Kaplan-Meier curve is shown that specifies the percentage of tumor-free mice against the weeks of age of the mice. Parental BALB NeuT mice developed tumors beginning around 14-15 weeks of age, with all mice developing tumors by 20 weeks of age. (BALBxPWK)F1 NeuT mice developed tumors beginning at 10 weeks of age and all mice were tumor positive by 15 weeks of age, significantly earlier than the parental BALB NeuT (*p* < 0.0001).

**Figure 3.**
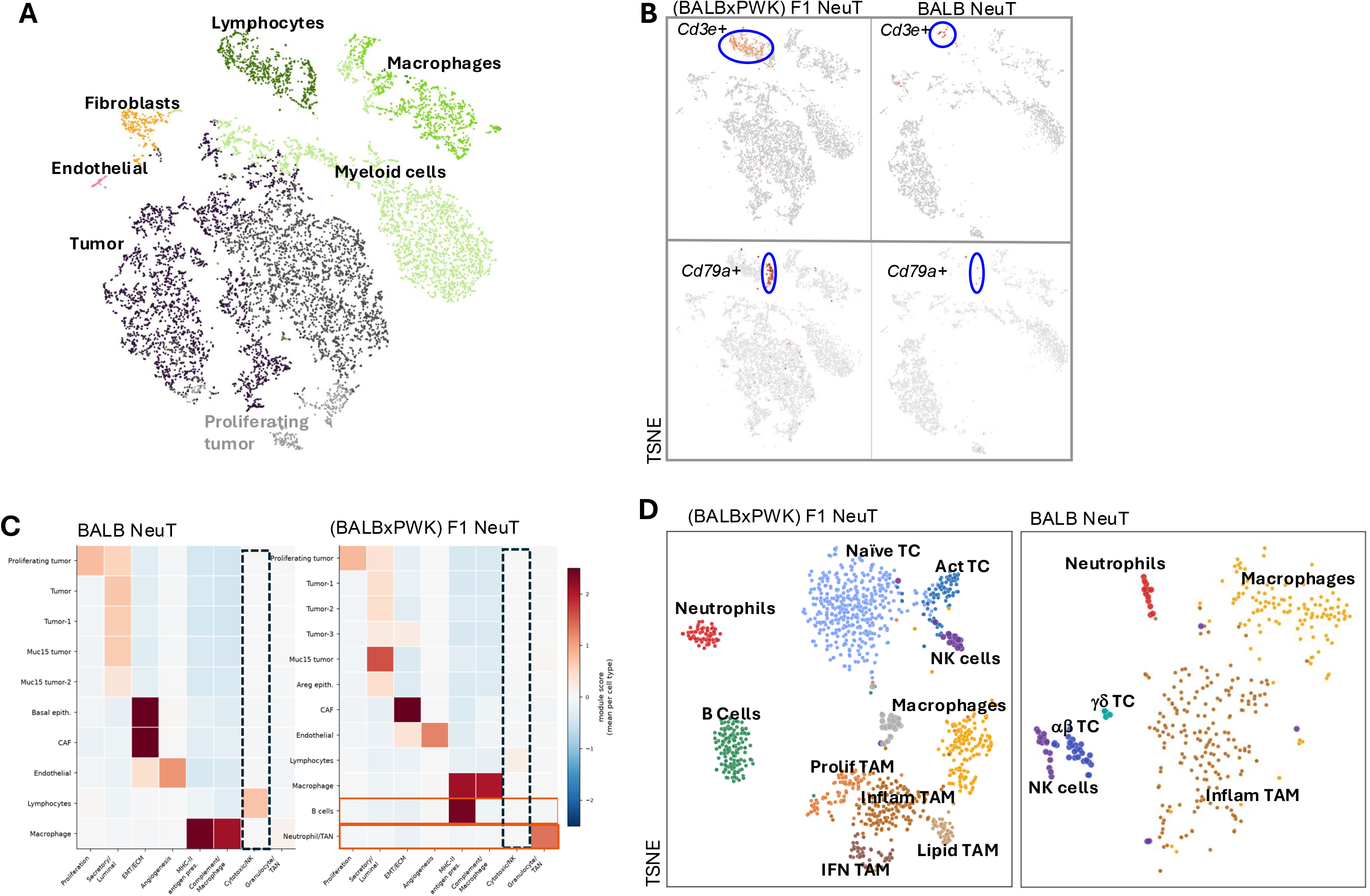
Single-cell RNA sequencing identified differences in lymphoid and myeloid representation between BALB NeuT and (BALBxPWK) F1 NeuT tumor datasets. **(A)** scRNA-Seq libraries from BALB NeuT and (BALBxPWK) F1 NeuT tumor tissue were aggregated and assessed for major cell types. Cell clustering identified tumor cells and proliferating tumor cells, fibroblasts, endothelial cells, lymphocytes, macrophages and myeloid cells. (B) Side-by-side comparison of the libraries showed greater frequency of *Cd3e*-expressing T cells and *Cd79a*-expressing B cells in the (BALBxPWK) F1 tumor tissue, despite the earlier tumor onset. **(C)** Comparison of the clusters showed that in addition to the a higher proportion of T cells and the presence of a B-cell population in the F1 NeuT tumors, there was also a distinct population of granulocytes/neutrophils that were enriched in the (BALBxPWK) F1 NeuT library but not detected in the BALB NeuT library. **(D)** t-SNE visualization of subpopulations within the immune-cell clusters.

To identify changes to the tumor microenvironment between the background strains, tumors from female BALB NeuT and (BALBxPWK) F1 NeuT mice were dissociated and analyzed by single-cell RNA sequencing (scRNA-Seq) (**Figure 3A**). Both datasets contained tumor epithelial, macrophage/myeloid, fibroblast, endothelial, and lymphoid populations. The F1 library contained a larger fraction of total immune cells than the BALB library (21.3% versus 13.8%; **Table 1**). While both libraries contained immune cell clusters (*Ptprc* or CD45), a major difference was found in lymphocyte representation (**Figure 3B**). *Cd3e*-expressing T cells made up the majority of the lymphocyte cluster for the (BALBxPWK) F1 NeuT sample and included a *Cd79a*-expressing B-cell population that was not detected in the BALB NeuT library (**Figure 3C**). To parse the differences further, immune cell populations were further sub-clustered (**Figure 3D**)(**Table 1**). In BALB NeuT mice, infiltrating immune cells were comprised of both αβ and γδ T cells, natural killer (NK) cells, a few neutrophils and substantial representation of monocyte-derived macrophages (macrophages) and tumor-associated macrophages (TAM). In the (BALBxPWK) F1 mixed background tumors, there was a large population of naïve T cells and TAM, with fewer overall macrophages, greater neutrophil representation, and a B-cell cluster not detected in BALB NeuT (**Figure 3D**, ***right***). In the mixed background, in addition to the naïve T cell cluster, there was also a distinct cluster of T cells expressing CD25/*Il2r.* Approximately 1/3^rd^ of these cells expressed *Foxp3* in addition to the canonical Treg markers *Ctla4*, *Tnfrsf4*, and *Ikzf2* while the remaining cells expressed *Il2r* together with *Cd8a* or *Cd4.* A closer inspection of the gene expression profiles within the lymphocyte clusters showed that cells in the BALB NeuT library expressed genes associated with cytotoxicity (*Tox, Ltk, Nkg7, Gzmb*) (**Supplemental Table 1**), which were lower or not detected in the mixed-background lymphocyte population. Together, the F1 lymphoid compartment showed greater representation of naïve/regulatory transcriptional states and lower expression of a cytotoxicity-associated gene program. It is important to note, however, that cytotoxic function was not directly measured.

**Table 1.**
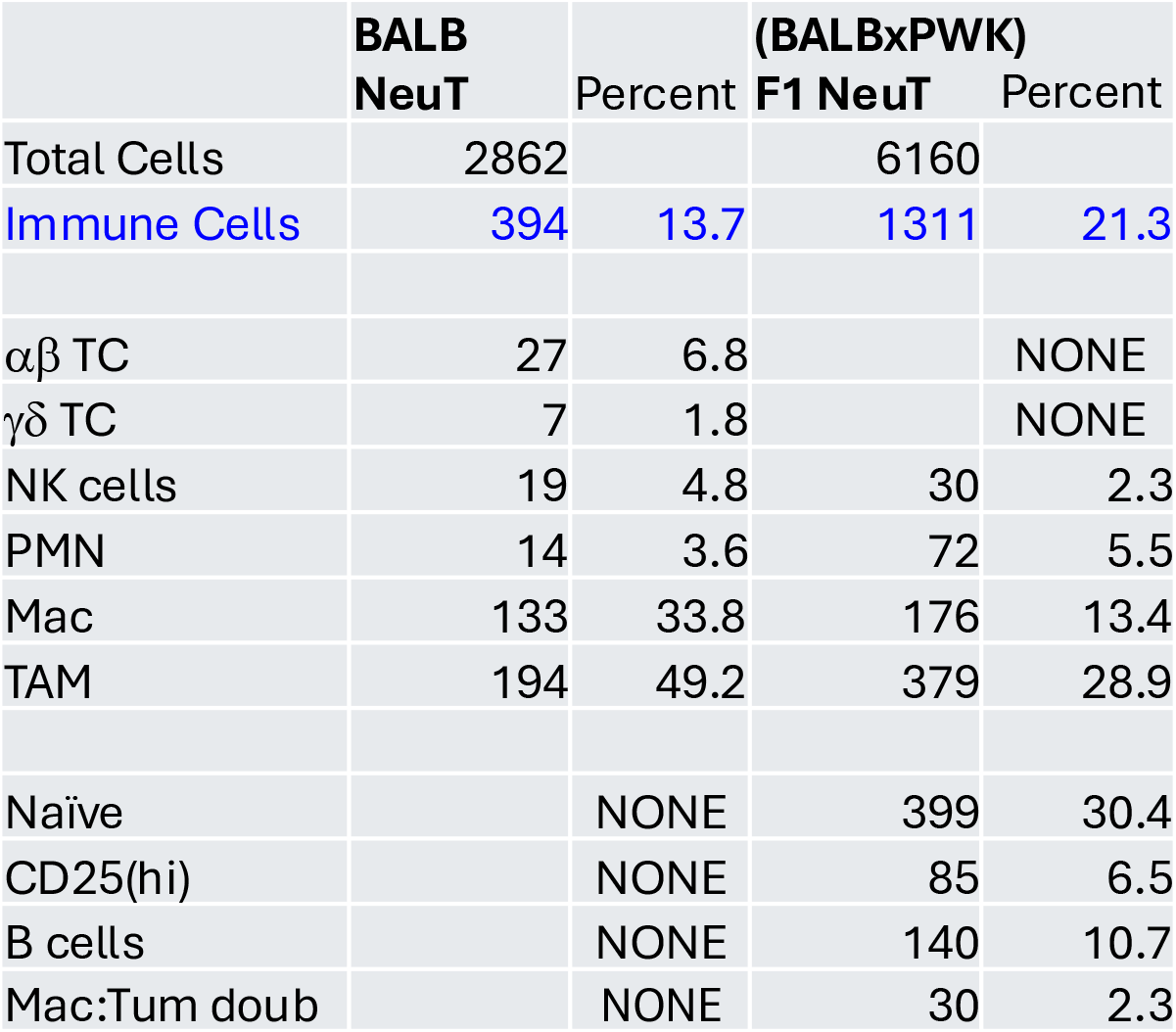
Immune cells identified in the single-cell RNA sequencing data.

|  | BALB<br>NeuT | Percent | (BALBxPWK)<br>F1 NeuT | Percent |
| --- | --- | --- | --- | --- |
| Total Cells | 2862 |  | 6160 |  |
| Immune Cells | 394 | 13.7 | 1311 | 21.3 |
| $\alpha\beta$ TC | 27 | 6.8 | | NONE |
| $\gamma\delta$ TC | 7 | 1.8 | | NONE |
| NK cells | 19 | 4.8 | 30 | 2.3 |
| PMN | 14 | 3.6 | 72 | 5.5 |
| Mac | 133 | 33.8 | 176 | 13.4 |
| TAM | 194 | 49.2 | 379 | 28.9 |
| Naïve |  | NONE | 399 | 30.4 |
| CD25(hi) |  | NONE | 85 | 6.5 |
| B cells |  | NONE | 140 | 10.7 |
| Mac:Tum doub |  | NONE | 30 | 2.3 |

Formalin-fixed paraffin-embedded (FFPE) blocks from BALB NeuT (n=5) and (BALBxPWK) F1 NeuT (n=5) tumors were assessed for protein abundance using data-independent acquisition (DIA). Forty-six proteins met the differential-abundance criteria, with an absolute log₂ fold change ≥ 1 and adjusted *p* value ≤ 0.05; 25 proteins were higher and 21 were lower in F1 tumors. Proteins with higher abundance included Fetub, Serpina1d, Papln, Fbln5, and Abcg2 (**Figure 4A, B**). Higher abundance of Fbln5 (elastic-fiber organization) and Papln (basement-membrane glycoprotein) was consistent with extracellular matrix (ECM) remodeling, while Fetub and Serpina1d are associated with protease regulation. Abcg2 is a drug efflux transporter and has been previously characterized as a marker of stem-like cells. Proteins with lower abundance included antigen-presentation components, including Stat1, Tapbp, Parp9, Dtx3l, and MHC class I and II proteins (**Figure 4B, C**). Although Tap1 and Tap2 did not meet the adjusted *p* < 0.05 threshold, their abundances were 1.5– and 1.4-fold lower, respectively (*p* = 0.06). Reactome analysis of the differentially abundant proteins identified reduced interferon-response and antigen-presentation pathways (**Figure 4D**).

**Figure 4.**
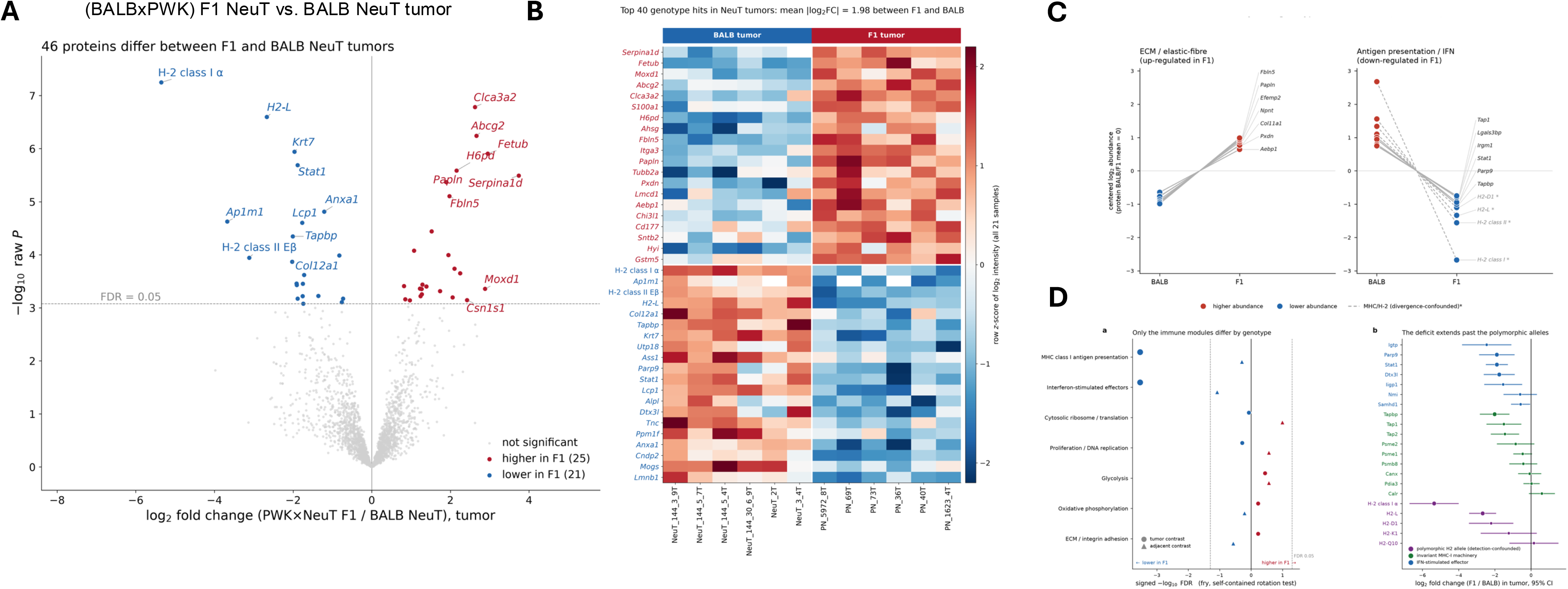
Proteomic analysis identified reduced MHC expression in (BALBxPWK) F1 NeuT tumors compared with BALB NeuT. FFPE tumor tissue from n=5 BALB NeuT and n=5 (BALBxPWK) F1 NeuT mice were submitted for data independent proteomic analysis (DIA). **(A)** A volcano plot showing proteins with the largest abundance differences in the (BALBxPWK) F1 NeuT mice compared with the BALB NeuT mice. **(B)** A heatmap of the top genotype-associated protein abundance differences across tumor samples. **(C)** Selected proteins with higher abundance in F1 tumors are associated with ECM and elastic-fiber biology, whereas selected proteins with lower abundance include antigen-presentation and interferon-response components. Points show replicate-level protein abundance values. (**D**) Pathway-level contrasts and selected protein effect estimates show reduced antigen-presentation and interferon-response programs in F1 tumors, including changes that extend beyond the polymorphic MHC proteins.

**Figure 5.**
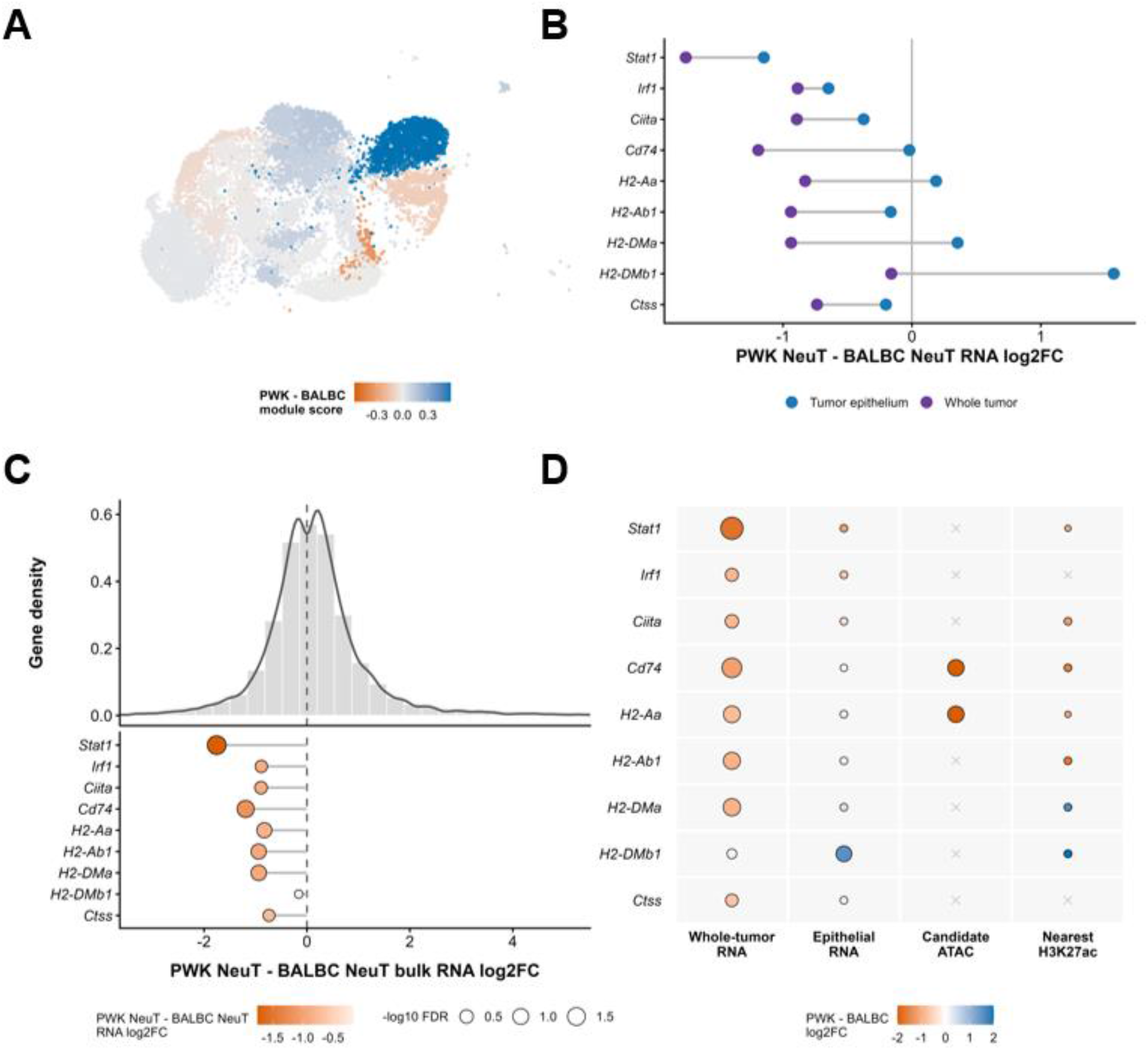
Focused cross-layer analysis of MHC II-associated regulation in BALB NeuT and (BALBxPWK) F1 NeuT tumors. **(A)** Replicate-averaged (BALBxPWK) F1 NeuT-minus-BALB NeuT differences in the seven-gene MHC II RNA module projected onto annotated epithelial states for descriptive visualization. **(B)** Comparison of whole-tumor bulk RNA-seq and epithelial multiome pseudobulk effects for *Stat1*/MHC II-associated genes. Values are log₂ fold changes for (BALBxPWK) F1 NeuT relative to BALB NeuT. **(C)** Whole-tumor bulk RNA-seq effect-size distribution for the nine-gene Stat1/MHC II-associated set compared with all remaining genes with finite estimates; the gene-set comparison is exploratory. (**D**) Cross-layer comparison of MHC II-associated genes across whole-tumor bulk RNA, epithelial multiome RNA, selected epithelial ATAC regions, and the nearest gene-associated bulk H3K27ac interval. Color indicates log₂ fold change and point size indicates statistical support where available.

To assess chromatin and transcriptional regulation of the antigen-presentation phenotype, we performed a focused analysis of companion multiome, bulk RNA-seq, and H3K27ac datasets (Geng et al., Submitted) with an emphasis on MHC II. Replicate-averaged MHC II module differences were projected onto the epithelial UMAP (**Figure 5A**). This analysis showed that MHC II module differences were heterogeneous across the annotated epithelial states, as opposed to a uniform reduction throughout the epithelial compartment. Reduced MHC II-associated expression effects were more pronounced in whole-tumor RNA profiles than in epithelial pseudobulk profiles (**Figure 5B**). However, epithelial effects were gene-dependent: the log₂ fold-change estimate for *Cd74*, the invariant chain of MHC II, was near zero, whereas estimates for *H2-Aa* and *H2-DMa* were modestly positive. An exploratory set of nine Stat1/MHC II-associated genes was evaluated for a distributional comparison. This gene set was shifted toward lower whole-tumor expression in (BALBxPWK) F1 NeuT tumors relative to the remaining genes with finite expression-effect estimates (one-sided Wilcoxon rank-sum test, nominal *p* = 3.1 × 10⁻⁵) (**Figure 5C**). This distributional shift extended beyond the individually significant decreases in *Stat1* and *Cd74* and was consistent with attenuation of the selected MHC II-associated program at the whole-tumor level. Finally, cross-layer comparison showed reduced epithelial chromatin accessibility at selected candidate regulatory regions near both *Cd74* and *H2-Aa* in (BALBxPWK) F1 NeuT tumors (**Figure 5D**). Together, these findings support whole-tumor attenuation of MHC II-associated transcription alongside heterogeneous epithelial expression and reduced accessibility at selected epithelial loci, rather than uniform epithelial suppression.

Together, the proteomic, RNA, and chromatin-accessibility analyses are consistent with reduced antigen-presentation capacity in (BALBxPWK) F1 NeuT tumors, alongside increased abundance of several ECM-related proteins.

## DISCUSSION

Using a genetically diverse F1 cross, we show that introducing the wild-derived PWK/PhJ background onto the BALB NeuT model of ErbB2/HER2-driven mammary carcinogenesis accelerates tumor onset by approximately four weeks without altering the kinetics of tumor growth once tumors are established, consistent with a quantitative trait locus (QTL) architecture in which distinct loci on chromosomes 1, X, and 10 are associated with onset and growth rate, respectively. This acceleration was accompanied by altered immune-cell representation in the scRNA-seq dataset: (BALBxPWK) F1 NeuT tumors showed greater representation of *Cd3e*-expressing T cells, a *Cd79a*-expressing B-cell population, and a distinct tumor-associated neutrophil cluster that were not detected or were less represented in BALB NeuT tumors. These compositional differences did not coincide with a stronger cytotoxic gene-expression program. Across complementary proteomic, transcriptomic, and chromatin-accessibility analyses, (BALBxPWK) F1 NeuT tumors showed coordinate down-regulation of a *Stat1*-driven antigen-presentation program. The combination of greater lymphocyte representation with reduced antigen-presentation markers is consistent with a tumor microenvironment in which immune-cell presence is uncoupled from robust antigen-presentation activity. This pattern departs from the relationship between MHC class I loss and lymphocyte infiltration reported in human epithelial cancers. In pancreatic ductal adenocarcinoma, tumors with reduced or absent HLA class I expression contained fewer infiltrating cytotoxic T cells and were associated with shorter survival, consistent with a model in which antigen presentation is required to recruit and sustain a cytotoxic T-cell response^51^.

In the (BALBxPWK) F1 NeuT tumors described here, T-cell representation showed the opposite pattern: despite lower MHC I/II and *Stat1* pathway protein abundance, T cells were more highly represented in the dissociated F1 scRNA-seq library than in the BALB NeuT library. However, these T cells showed reduced or undetected expression of the cytotoxicity-associated genes *Tox, Ltk, Nkg7*, and *Gzmb* that were present in BALB NeuT tumor-infiltrating lymphocytes, and sub-clustering resolved the F1 lymphoid compartment into naive and CD25(hi)/*Il2r*-expressing populations, roughly one-third of which expressed *Foxp3* together with *Ctla4, Tnfrsf4*, and *Ikzf2*, a phenotype more consistent with a naive/regulatory state than with an activated effector response. These transcriptional data do not establish impaired cytotoxic function, but they support a model in which lymphocyte representation and acquisition of an effector program can be dissociated.

The coordinate loss of *Stat1, Tapbp, Parp9*, and *Dtx3l* alongside MHC class I and II proteins is consistent with a shared upstream regulatory node rather than independent, unrelated changes. Stat1, together with IRF-1, is required for interferon-driven transactivation of the *Tap1* and *Lmp2* (*Psmb9*) genes that supply peptides to the MHC class I loading complex^52^, and Stat1-dependent signaling through the interferon-stimulated response element is likewise necessary for full HLA class I transactivation^53^. Parp9 and Dtx3l function as an obligate heterodimer downstream of Stat1 that amplifies interferon-stimulated gene expression, including components of the antigen-presentation pathway^54^. The parallel, sub-significant reduction in *Tap1* and *Tap2* (1.5– and 1.4-fold, *p* = 0.06) is directionally consistent with this model, and pathway-level testing of the proteomic data confirmed that MHC class I antigen presentation and interferon-stimulated effector modules were the most significantly down-regulated gene sets distinguishing the two genotypes, rather than being driven by one or two outlier proteins. Together, these results are consistent with reduced STAT1-associated signaling in (BALBxPWK) F1 NeuT tumors, accompanied by coordinated decreases in MHC I/II components, Tapbp, and the Parp9-Dtx3l module.

The multiome data extend this *Stat1*-centered model to MHC class II regulation and to chromatin accessibility. CIITA, the master transactivator of MHC class II and the invariant-chain gene *Cd74*, is itself an interferon-inducible gene that requires cooperative Stat1 and USF-1 binding for full induction^55^, and CIITA deficiency is sufficient to abolish MHC class II expression^56^. Consistent with this hierarchy, *Stat1, Irf1*, and *Ciita* were each reduced in (BALBxPWK) F1 NeuT whole-tumor RNA, and this reduction extended downstream to *Cd74* and to chromatin accessibility at candidate regulatory regions near *Cd74* and *H2-Aa*. Notably, CD74 was not among the 46 differentially abundant proteins identified by proteomics; support for its down-regulation comes from whole-tumor bulk RNA and chromatin-accessibility analyses rather than differential protein abundance. The lack of a proteomic difference could reflect assay sensitivity, protein turnover, or incomplete protein coverage. The multiome data also showed that this attenuation was not uniform across the tumor: *H2-DMa* and *H2-DMb1* had positive epithelial RNA and/or chromatin-accessibility effect estimates even as whole-tumor RNA effects were reduced. This heterogeneity indicates that the antigen-presentation signal likely reflects differences in cellular composition and cell state, together with gene-specific epithelial effects, rather than uniform cell-intrinsic suppression of MHC II throughout the tumor epithelium.

A specific concern for proteomic comparison across genetically divergent strains is that classical MHC class I and II genes are highly polymorphic. Strain-specific sequence differences can alter tryptic peptide composition, peptide mapping, and quantification against a reference proteome, potentially contributing to apparent abundance differences between BALB/c– and PWK/PhJ-derived H2 haplotypes. Several of the largest-magnitude changes in the proteomic dataset involved these polymorphic loci. However, the same direction of change was also observed for non-MHC pathway members, including *Stat1, Tapbp*, and *Parp9*, arguing that the pattern extends beyond the most polymorphic H2 loci and is not explained solely by MHC haplotype divergence. Directionally concordant changes were also observed in tumor-adjacent tissue for selected proteins, raising the possibility that part of this phenotype precedes or accompanies tumor formation; this observation is exploratory and requires dedicated normal-tissue comparisons.

In parallel with reduced antigen presentation, (BALBxPWK) F1 NeuT tumors up-regulated a distinct set of proteins involved in extracellular-matrix and elastic-fiber organization (*Fbln5, Papln, Fetub*, *Serpina1d*) alongside MET-PTK2 signaling activity, a pathway associated with tumor cell motility. *Abcg2*, a canonical target of the oxidative-stress-responsive transcription factor NFE2L2 (Nrf2), was among the most strongly up-regulated proteins and has been characterized elsewhere as a marker of stem-like tumor cells and a mediator of xenobiotic efflux. Whether ECM remodeling and the antigen-presentation-low phenotype are mechanistically linked, share a common upstream driver, or represent independent consequences of the same genetic background cannot be resolved from these data. The present results support parallel effects on matrix remodeling and immune recognition without establishing a direct mechanistic connection between them.

Future work should prioritize fine-mapping of the chromosome 1, X, and 10 QTL intervals to identify causal genes, functional validation of *Cd74* and CIITA regulation at the protein level, and direct cytotoxicity assays of tumor-infiltrating T cells to determine whether restoring antigen presentation is sufficient to license an effector response in the (BALBxPWK) F1 NeuT background. Because the phenotype described here combines greater lymphocyte representation with reduced antigen-presentation markers, it may also be informative to test how tumors with this genetic background respond to immune-checkpoint blockade or HER2-targeted therapy relative to BALB NeuT tumors, since immune-infiltrated but antigen-presentation-low tumors may respond differently to these interventions than either immune-excluded or fully immunocompetent tumors.

## CONCLUSION

Together, these findings show that introduction of the PWK/PhJ wild-derived genetic background associates earlier tumor onset with reduced antigen-presentation activity; even in the face of greater lymphocyte presence. (BALBxPWK) F1 NeuT tumors showed coordinated reduction of STAT1-associated antigen-presentation programs across proteomic, transcriptomic, and chromatin analyses, together with a lymphoid compartment enriched for naïve and regulatory transcriptional states as opposed to a strong cytotoxic program. These results identify host genetic background as an important modifier of the relationship between tumor development and antitumor immunity and provide a framework for testing how inherited variation influences response to HER2-targeted and immune-based therapies.

## AUTHOR CONTRIBUTIONS

<u>Conception</u>. L.P., B.L.K. and J.B.J

<u>Methodology</u>. B.L.K. and J.B.J

<u>Acquisition</u>. K.W., R.R.R., B.L.K. and J.B.J

Software. J.B.J.

<u>Validation</u>. R.R.R. and J.B.J.

<u>Formal Analysis</u>. R.R.R., B.L.K. and J.B.J

<u>Investigation</u>. R.R.R., R.G., B.L.K. and J.B.J

<u>Resources</u>. B.L.K. and J.B.J.

<u>Data Curation</u>. R.R.R, B.L.K. and J.B.J

Writing – original draft preparation. J.B.J

Writing – Review and editing. B.L.K. and J.B.J

<u>Visualization</u>. B.L.K. and J.B.J.

<u>Supervision</u>. B.L.K and J.B.J

Project Administration. J.B.J.

<u>Funding Acquisition</u>. B.L.K. and J.B.J.

## INSTITUTIONAL REVIEW BOARD STATEMENT

All procedures were performed in accordance with the US Public Health Service Policy on the Use of Laboratory Animals and with approval by Michigan State University’s Institutional Animal Care and Use Committee.

## DATA AVAILABILITY STATEMENT

The authors have deposited the raw data that was used to analyze single cell RNA sequencing (scRNA-Seq), proteomics data and multiome data to the NCBI Gene Expression Omnibus (GEO).

## ACKNOWLEDGMENTS

The authors would like to acknowledge the IDeA Proteomics Core (Little Rock, AR), the Center for Molecular Medicine and Genetics (Wayne State University, Detroit, MI) and the Genetics and Genomics Core at Michigan State University (East Lansing, MI) for their assistance with data collection. The authors would also like to acknowledge the animal care staff at both Wayne State University and Michigan State University for their excellent care of the research animals.

## REFERENCES

1. Bose R, Kavuri SM, Searleman AC, et al. Activating HER2 mutations in HER2 gene amplification negative breast cancer. Cancer Discov. Feb 2013;3(2):224–37. doi:10.1158/2159-8290.CD-12-0349

2. Palmieri D, Bronder JL, Herring JM, et al. Her-2 overexpression increases the metastatic outgrowth of breast cancer cells in the brain. Cancer Res. May 1 2007;67(9):4190–8. doi:10.1158/0008-5472.CAN-06-3316

3. Cheng X, Sun Y, Highkin M, et al. Breast Cancer Mutations HER2V777L and PIK3CAH1047R Activate the p21-CDK4/6-Cyclin D1 Axis to Drive Tumorigenesis and Drug Resistance. Cancer Res. Sep 1 2023;83(17):2839–2857. doi:10.1158/0008-5472.CAN-22-3558

4. Abbasvandi F, Bayat M, Akbari A, et al. Tumor characteristics and survival rate of HER2-low breast cancer patients: a retrospective cohort study. Sci Rep. Oct 4 2023;13(1):16719. doi:10.1038/s41598-023-43186-8

5. Rovero S, Amici A, Di Carlo E, et al. DNA vaccination against rat her-2/Neu p185 more effectively inhibits carcinogenesis than transplantable carcinomas in transgenic BALB/c mice. J Immunol. Nov 1 2000;165(9):5133–42. doi:10.4049/jimmunol.165.9.5133

6. Bargmann CI, Hung MC, Weinberg RA. Multiple independent activations of the neu oncogene by a point mutation altering the transmembrane domain of p185. Cell. Jun 6 1986;45(5):649–57. doi:10.1016/0092-8674(86)90779-8

7. Conti L, Ruiu R, Barutello G, et al. Microenvironment, oncoantigens, and antitumor vaccination: lessons learned from BALB-neuT mice. Biomed Res Int. 2014;2014:534969. doi:10.1155/2014/534969

8. Jacob J, Radkevich O, Forni G, et al. Activity of DNA vaccines encoding self or heterologous Her-2/neu in Her-2 or neu transgenic mice. Cell Immunol. Apr 2006;240(2):96–106. doi:10.1016/j.cellimm.2006.07.002

9. Jacob JB, Kong YC, Nalbantoglu I, Snower DP, Wei WZ. Tumor regression following DNA vaccination and regulatory T cell depletion in neu transgenic mice leads to an increased risk for autoimmunity. J Immunol. May 1 2009;182(9):5873–81. doi:10.4049/jimmunol.0804074

10. Jacob JB, Quaglino E, Radkevich-Brown O, et al. Combining human and rat sequences in her-2 DNA vaccines blunts immune tolerance and drives antitumor immunity. Cancer Res. Jan 1 2010;70(1):119–28. doi:10.1158/0008-5472.CAN-09-2554

11. Jacob JB, Wei KC, Bepler G, et al. Identification of actionable targets for breast cancer intervention using a diversity outbred mouse model. iScience. Apr 21 2023;26(4):106320. doi:10.1016/j.isci.2023.106320

12. Broman KW, Gatti DM, Simecek P, et al. R/qtl2: Software for Mapping Quantitative Trait Loci with High-Dimensional Data and Multiparent Populations. Genetics. Feb 2019;211(2):495–502. doi:10.1534/genetics.118.301595

13. Gatti DM, Svenson KL, Shabalin A, et al. Quantitative trait locus mapping methods for diversity outbred mice. G3 (Bethesda). Sep 18 2014;4(9):1623–33. doi:10.1534/g3.114.013748

14. Hanahan D. Hallmarks of Cancer: New Dimensions. Cancer Discov. Jan 2022;12(1):31–46. doi:10.1158/2159-8290.CD-21-1059

15. Hanahan D. Hallmarks of cancer-Then and now, and beyond. Cell. Apr 16 2026;189(8):2254–2277. doi:10.1016/j.cell.2025.12.049

16. Hanahan D, Weinberg RA. Hallmarks of cancer: the next generation. Cell. Mar 4 2011;144(5):646–74. doi:10.1016/j.cell.2011.02.013

17. Tao J, Li Y, Liu YQ, et al. Restoration of the expression of transports associated with antigen processing in human malignant melanoma increases tumor-specific immunity. J Invest Dermatol. Aug 2008;128(8):1991–6. doi:10.1038/jid.2008.10

18. Rodig SJ, Gusenleitner D, Jackson DG, et al. MHC proteins confer differential sensitivity to CTLA-4 and PD-1 blockade in untreated metastatic melanoma. Sci Transl Med. Jul 18 2018;10(450)doi:10.1126/scitranslmed.aar3342

19. Balko JM, Licata L, Wang XQ, et al. Tumor-specific major histocompatibility-II expression predicts pathological complete response to atezolizumab combined to chemotherapy in triple-negative breast cancer. NPJ Breast Cancer. Sep 29 2025;11(1):103. doi:10.1038/s41523-025-00828-6

20. Kono K, Sato E, Naganuma H, et al. Trastuzumab (Herceptin) enhances class I-restricted antigen presentation recognized by HER-2/neu-specific T cytotoxic lymphocytes. Clin Cancer Res. Apr 1 2004;10(7):2538–44. doi:10.1158/1078-0432.ccr-03-0424

21. Griguolo G, Pascual T, Dieci MV, Guarneri V, Prat A. Interaction of host immunity with HER2-targeted treatment and tumor heterogeneity in HER2-positive breast cancer. J Immunother Cancer. Mar 29 2019;7(1):90. doi:10.1186/s40425-019-0548-6

22. Dhatchinamoorthy K, Colbert JD, Rock KL. Cancer Immune Evasion Through Loss of MHC Class I Antigen Presentation. Front Immunol. 2021;12:636568. doi:10.3389/fimmu.2021.636568

23. Koebel CM, Vermi W, Swann JB, et al. Adaptive immunity maintains occult cancer in an equilibrium state. Nature. Dec 6 2007;450(7171):903–7. doi:10.1038/nature06309

24. Henle AM, Nassar A, Puglisi-Knutson D, Youssef B, Knutson KL. Downregulation of TAP1 and TAP2 in early stage breast cancer. PLoS One. 2017;12(11):e0187323. doi:10.1371/journal.pone.0187323

25. Ogino T, Shigyo H, Ishii H, et al. HLA class I antigen down-regulation in primary laryngeal squamous cell carcinoma lesions as a poor prognostic marker. Cancer Res. Sep 15 2006;66(18):9281–9. doi:10.1158/0008-5472.CAN-06-0488

26. Vitale M, Rezzani R, Rodella L, et al. HLA class I antigen and transporter associated with antigen processing (TAP1 and TAP2) down-regulation in high-grade primary breast carcinoma lesions. Cancer Res. Feb 15 1998;58(4):737–42.

27. Seliger B. Molecular mechanisms of MHC class I abnormalities and APM components in human tumors. Cancer Immunol Immunother. Nov 2008;57(11):1719–26. doi:10.1007/s00262-008-0515-4

28. Campoli M, Ferrone S. HLA antigen changes in malignant cells: epigenetic mechanisms and biologic significance. Oncogene. Oct 6 2008;27(45):5869–85. doi:10.1038/onc.2008.273

29. Pedersen MH, Hood BL, Beck HC, Conrads TP, Ditzel HJ, Leth-Larsen R. Downregulation of antigen presentation-associated pathway proteins is linked to poor outcome in triple-negative breast cancer patient tumors. Oncoimmunology. 2017;6(5):e1305531. doi:10.1080/2162402X.2017.1305531

30. Collins T, Korman AJ, Wake CT, et al. Immune interferon activates multiple class II major histocompatibility complex genes and the associated invariant chain gene in human endothelial cells and dermal fibroblasts. Proc Natl Acad Sci U S A. Aug 1984;81(15):4917–21. doi:10.1073/pnas.81.15.4917

31. Jabrane-Ferrat N, Faille A, Loiseau P, Poirier O, Charron D, Calvo F. Effect of gamma interferon on HLA class-I and –II transcription and protein expression in human breast adenocarcinoma cell lines. Int J Cancer. Jun 15 1990;45(6):1169–76. doi:10.1002/ijc.2910450630

32. da Silva GB, Silva TG, Duarte RA, et al. Expression of the Classical and Nonclassical HLA Molecules in Breast Cancer. Int J Breast Cancer. 2013;2013:250435. doi:10.1155/2013/250435

33. Feinmesser M, Sulkes A, Morgenstern S, Sulkes J, Stern S, Okon E. HLA-DR and beta 2 microglobulin expression in medullary and atypical medullary carcinoma of the breast: histopathologically similar but biologically distinct entities. J Clin Pathol. Apr 2000;53(4):286–91. doi:10.1136/jcp.53.4.286

34. Park IA, Hwang SH, Song IH, et al. Expression of the MHC class II in triple-negative breast cancer is associated with tumor-infiltrating lymphocytes and interferon signaling. PLoS One. 2017;12(8):e0182786. doi:10.1371/journal.pone.0182786

35. Forero A, Li Y, Chen D, et al. Expression of the MHC Class II Pathway in Triple-Negative Breast Cancer Tumor Cells Is Associated with a Good Prognosis and Infiltrating Lymphocytes. Cancer Immunol Res. May 2016;4(5):390–9. doi:10.1158/2326-6066.CIR-15-0243

36. Lee HJ, Song IH, Park IA, et al. Differential expression of major histocompatibility complex class I in subtypes of breast cancer is associated with estrogen receptor and interferon signaling. Oncotarget. May 24 2016;7(21):30119–32. doi:10.18632/oncotarget.8798

37. Mimura K, Ando T, Poschke I, et al. T cell recognition of HLA-A2 restricted tumor antigens is impaired by the oncogene HER2. Int J Cancer. Jan 15 2011;128(2):390–401. doi:10.1002/ijc.25613

38. Restifo NP, Marincola FM, Kawakami Y, Taubenberger J, Yannelli JR, Rosenberg SA. Loss of functional beta 2-microglobulin in metastatic melanomas from five patients receiving immunotherapy. J Natl Cancer Inst. Jan 17 1996;88(2):100–8. doi:10.1093/jnci/88.2.100

39. Herrmann F, Lehr HA, Drexler I, et al. HER-2/neu-mediated regulation of components of the MHC class I antigen-processing pathway. Cancer Res. Jan 1 2004;64(1):215–20. doi:10.1158/0008-5472.can-2522-2

40. Inoue M, Mimura K, Izawa S, et al. Expression of MHC Class I on breast cancer cells correlates inversely with HER2 expression. Oncoimmunology. Oct 1 2012;1(7):1104–1110. doi:10.4161/onci.21056

41. Fang Y, Wang L, Wan C, et al. MAL2 drives immune evasion in breast cancer by suppressing tumor antigen presentation. J Clin Invest. Jan 4 2021;131(1)doi:10.1172/JCI140837

42. Zhao MZ, Sun Y, Jiang XF, Liu L, Liu L, Sun LX. Promotion on NLRC5 upregulating MHC-I expression by IFN-gamma in MHC-I-deficient breast cancer cells. Immunol Res. Dec 2019;67(6):497–504. doi:10.1007/s12026-019-09111-w

43. Boggio K, Nicoletti G, Di Carlo E, et al. Interleukin 12-mediated prevention of spontaneous mammary adenocarcinomas in two lines of Her-2/neu transgenic mice. J Exp Med. Aug 3 1998;188(3):589–96. doi:10.1084/jem.188.3.589

44. Lucchini F, Sacco MG, Hu N, et al. Early and multifocal tumors in breast, salivary, harderian and epididymal tissues developed in MMTY-Neu transgenic mice. Cancer Lett. Jul 10 1992;64(3):203–9. doi:10.1016/0304-3835(92)90044-v

45. Ray R, Wei WZ, Jacob J. Dissociation of Murine Mammary Gland and Tumor Tissues to Create Highly Viable Single-Cell Suspensions. J Vis Exp. Aug 21 2026;(234)doi:10.3791/72844

46. Zhou L, Adrianto I, Wang J, Wu X, Datta I, Mi QS. Single-Cell RNA-Seq Analysis Uncovers Distinct Functional Human NKT Cell Sub-Populations in Peripheral Blood. Front Cell Dev Biol. 2020;8:384. doi:10.3389/fcell.2020.00384

47. Dorrity MW, Saunders LM, Queitsch C, Fields S, Trapnell C. Dimensionality reduction by UMAP to visualize physical and genetic interactions. Nat Commun. Mar 24 2020;11(1):1537. doi:10.1038/s41467-020-15351-4

48. Xie Z, Bailey A, Kuleshov MV, et al. Gene Set Knowledge Discovery with Enrichr. Curr Protoc. Mar 2021;1(3):e90. doi:10.1002/cpz1.90

49. Tabula Muris C, Overall c, Logistical c, et al. Single-cell transcriptomics of 20 mouse organs creates a Tabula Muris. Nature. Oct 2018;562(7727):367–372. doi:10.1038/s41586-018-0590-4

50. Geng R, Ray R, Jacob J, Kidder BL. PWK genetic background reprograms heterochromatin and malignant epithelial states in Neu-driven mammary tumors. BioRxiv: BioRxiv; 2026.

51. Ryschich E, Notzel T, Hinz U, et al. Control of T-cell-mediated immune response by HLA class I in human pancreatic carcinoma. Clin Cancer Res. Jan 15 2005;11(2 Pt 1):498–504.

52. Brucet M, Marques L, Sebastian C, Lloberas J, Celada A. Regulation of murine Tap1 and Lmp2 genes in macrophages by interferon gamma is mediated by STAT1 and IRF-1. Genes Immun. Jan 2004;5(1):26–35. doi:10.1038/sj.gene.6364035

53. Gobin SJ, van Zutphen M, Woltman AM, van den Elsen PJ. Transactivation of classical and nonclassical HLA class I genes through the IFN-stimulated response element. J Immunol. Aug 1 1999;163(3):1428–34.

54. Zhang Y, Mao D, Roswit WT, et al. PARP9-DTX3L ubiquitin ligase targets host histone H2BJ and viral 3C protease to enhance interferon signaling and control viral infection. Nat Immunol. Dec 2015;16(12):1215–27. doi:10.1038/ni.3279

55. Muhlethaler-Mottet A, Di Berardino W, Otten LA, Mach B. Activation of the MHC class II transactivator CIITA by interferon-gamma requires cooperative interaction between Stat1 and USF-1. Immunity. Feb 1998;8(2):157–66. doi:10.1016/s1074-7613(00)80468-9

56. Steimle V, Siegrist CA, Mottet A, Lisowska-Grospierre B, Mach B. Regulation of MHC class II expression by interferon-gamma mediated by the transactivator gene CIITA. Science. Jul 1 1994;265(5168):106–9. doi:10.1126/science.8016643

