## Supplemental Information for "Early-onset spontaneous murine mammary tumors show increased lymphocyte representation and reduced MHC antigen presentation"

### Jacob-Kidder_Supplemental Data

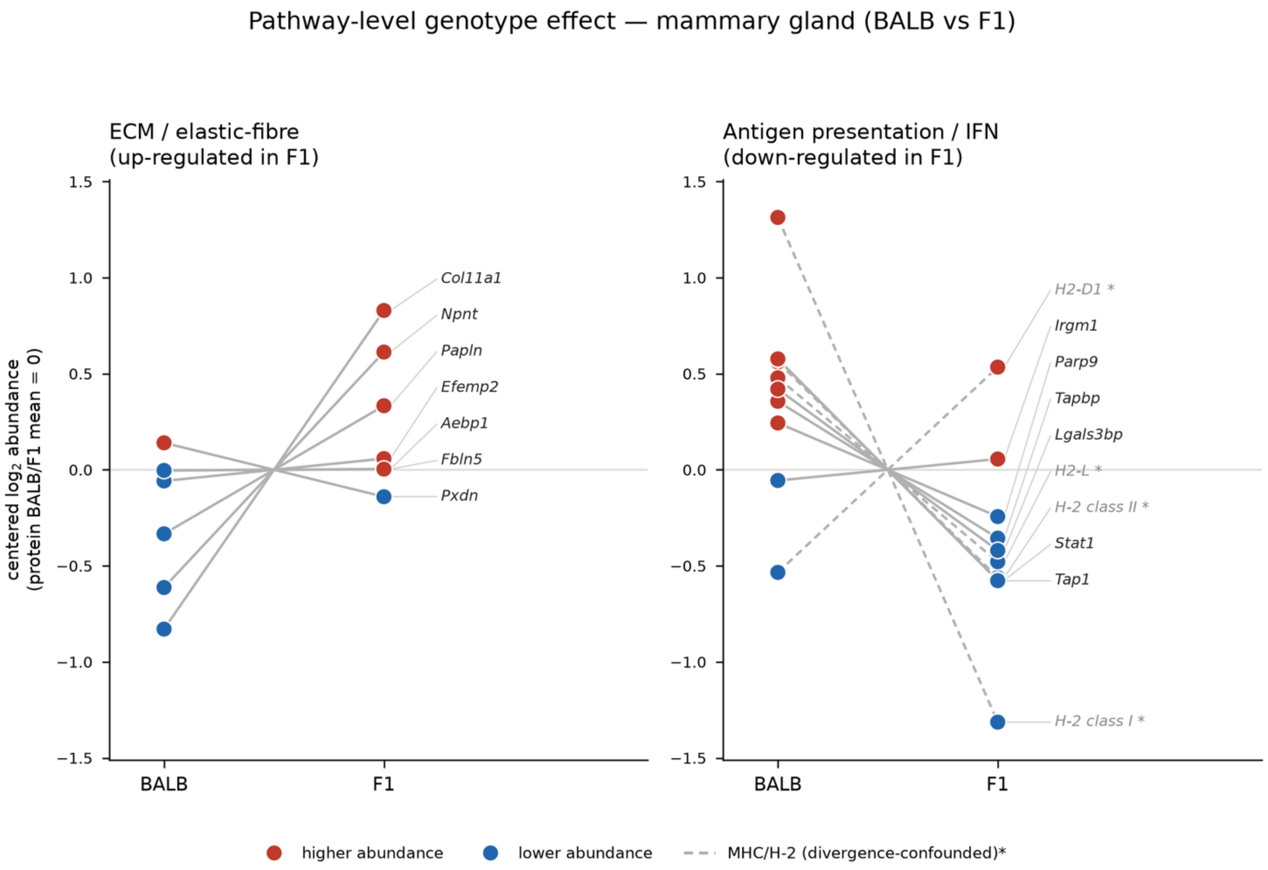


**Supplemental Figure 1.** Tumor-adjacent concordance of up-regulated ECM proteins and down-regulated antigen presentation proteins

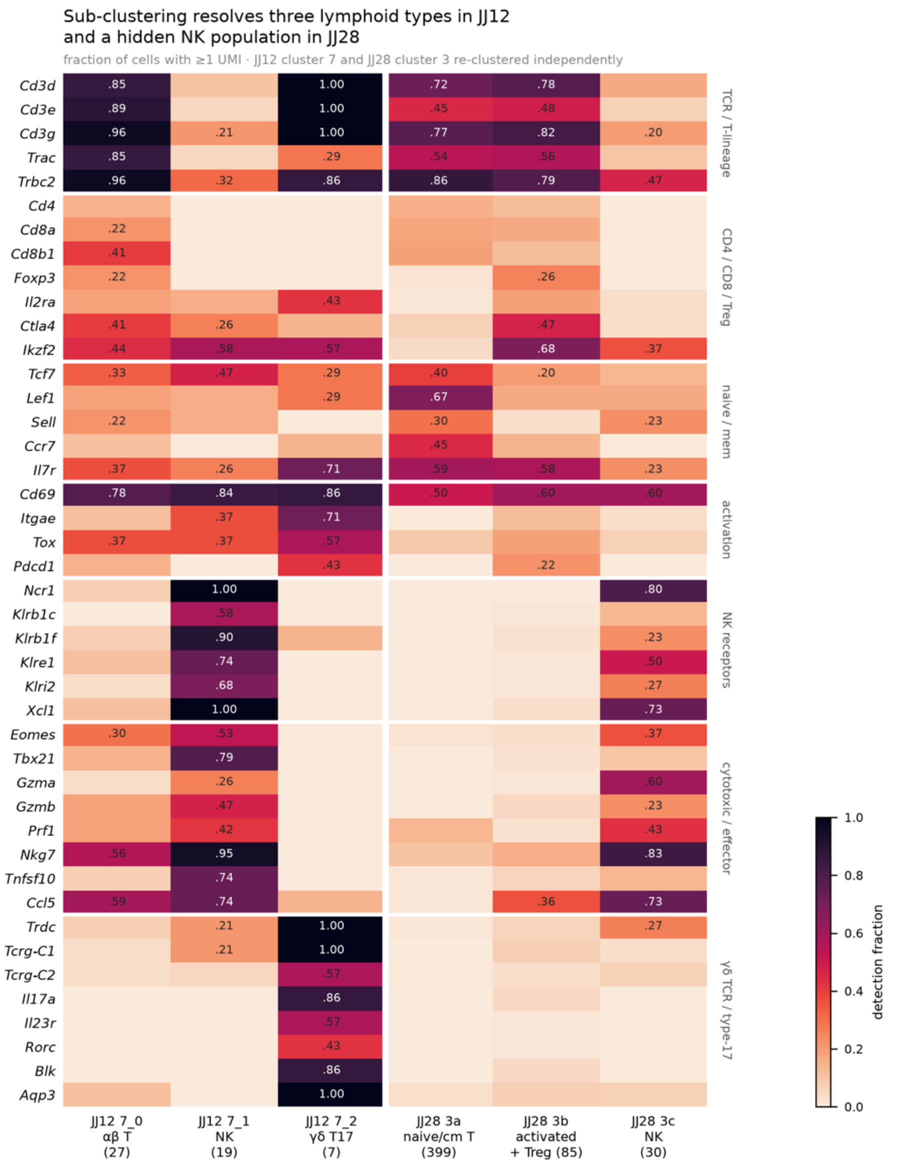


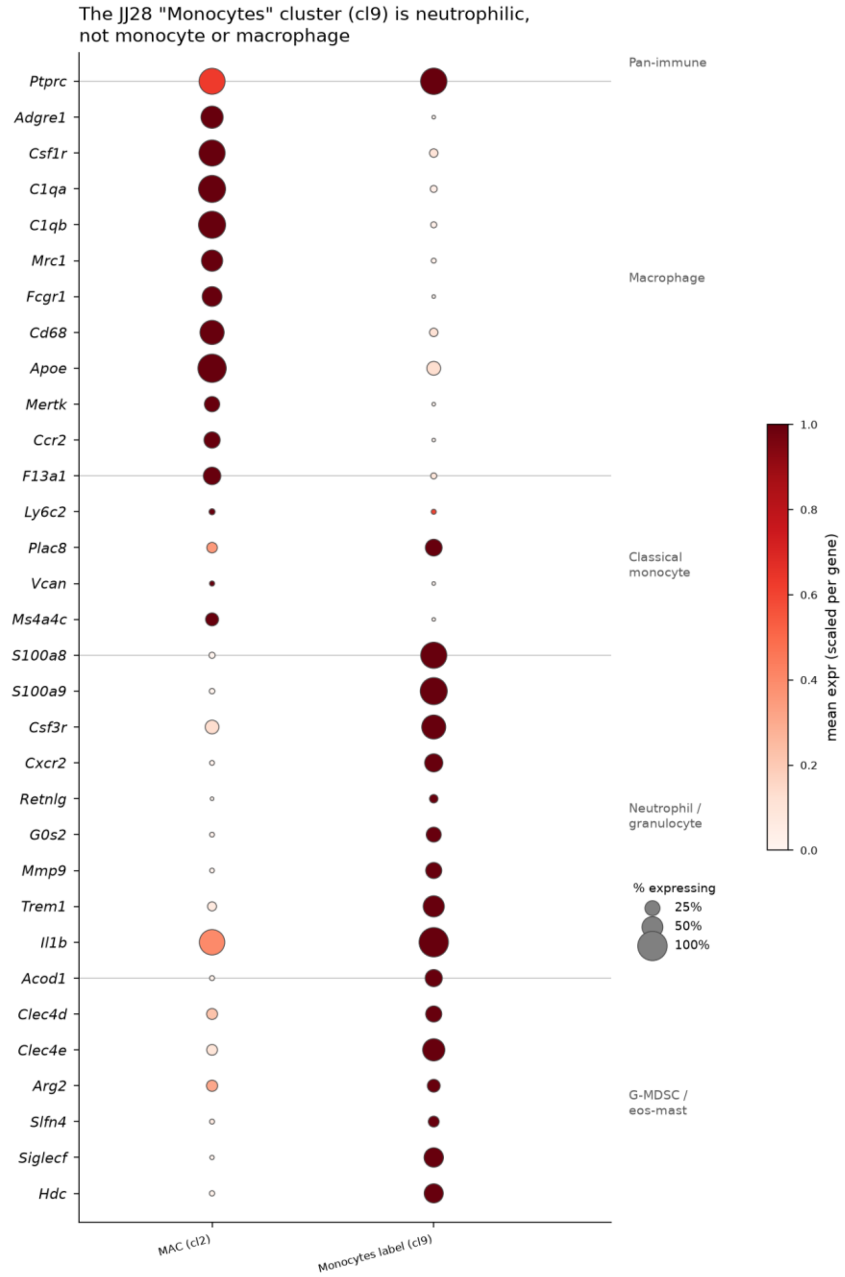


Supplemental Figure 2.

Supplemental Figure 3. Pathway enrichment

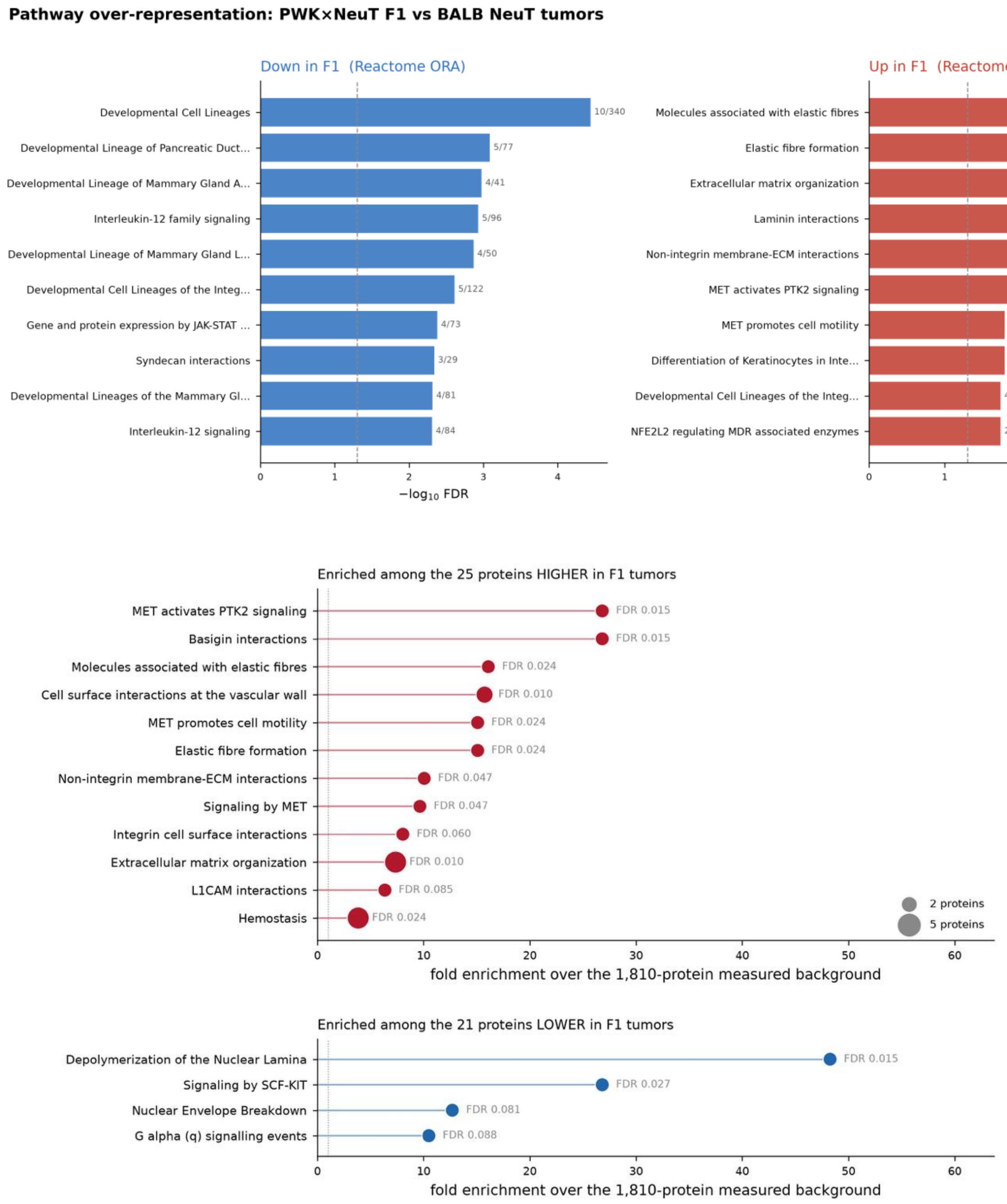
